# Tools of Hibernation Measurement and Interpretation (TOHMIN) for quantifying various values from body temperature fluctuation during hibernation

**DOI:** 10.1101/2024.12.09.626892

**Authors:** Reo Otsuka, Yutaro Shimoyama, Satoshi Nakagawa, Yoshifumi Yamaguchi

## Abstract

Hibernation is a fascinating physiological phenomenon that dramatically reduces basal metabolism and thermogenesis, resulting in a large deviation in body temperature (Tb) from homeothermic ranges in mammals. Although high-resolution long-term Tb recording in wild or laboratory animals has become possible through data loggers, few standardized methods to analyze details of hibernation patterns are available, making it difficult to reproduce and compare the results across different studies and species. To facilitate the analysis of hibernation patterns and accelerate hibernation research, we developed an open-source program, tools of hibernation measurement and interpretation (TOHMIN). As a proof of concept, we analyzed a dataset from two pilot studies on (1) the effects of distinct diets on hibernation patterns and (2) differences in hibernation patterns between males and females in a mammalian hibernator, Syrian hamster (*Mesocricetus auratus*), and found previously undetectable fine-scale differences in hibernation patterns. First, different types of diets affected the duration of periodic arousal. Second, females maintained higher body temperatures during periodic arousal than males. Third, the duration of the pre-hibernation period was negatively correlated with the hibernation period for this species. Thus, TOHMIN accelerates studies to assess the effects of various experimental manipulations on hibernation phenotypes in mammals.

## 1. Introduction

Hibernation is an adaptive strategy to endure seasons with little water or food by suppressing metabolism and thermogenesis, thereby being low body temperature. This remarkable adaptation has been observed in various mammalian species, ranging from small rodents to bears, and has fascinated researchers for many decades (1, 2); small mammalian hibernators, including ground squirrels, dormice, hamsters, and bats, are known to spend months for hibernation, alternating between two dramatically different phases: deep torpor (DT) and periodic (interbout) arousal (PA) (3–6). In DT, the metabolic rate, heart rate, and respiratory rate are remarkably suppressed, and Tb approaches the ambient temperature, with the animal remaining in an immobile state for several days to several weeks (7). The duration of DT varies by species and ambient temperature, reflecting the diverse ecological niches occupied by hibernators (8). The DT phase is interrupted by PA, which occurs spontaneously and is achieved via non-shivering thermogenesis in brown adipose tissue and shivering thermogenesis in skeletal muscle, allowing the animal to rewarm to normothermia (9). Rewarming from DT to normothermic PA requires considerable energy, which highlights the physiological costs associated with this process (10). The normothermic PA phase lasts for hours to days. During PA, food-storing hibernators, such as hamsters and chipmunks, eat stored food, whereas fat-storing hibernators do not, reflecting the different strategies for energy management during hibernation (11, 12).

The physiological, neural, and molecular mechanisms that regulate hibernation remain unclear. While circannual rhythms and their interactions with environmental cues play a crucial role in the regulation of hibernation in seasonal hibernators (13), many questions remain regarding the precise regulation of torpor and arousal. Central to these questions is how hibernators achieve and maintain metabolic suppression at low body temperature. Recent studies have shown that the degree and pattern of metabolic suppression can vary significantly among individuals and across species (8). This variability suggests that factors beyond circannual rhythms may influence hibernation strategies. Of particular interest are the potential roles of nutrition and sex in modulating hibernation patterns. For instance, pre-hibernation feeding behaviors and dietary composition have been shown to affect lipid storage and utilization during torpor (14). Moreover, emerging evidence indicates that male and female hibernators may employ different metabolic strategies during hibernation, possibly because of differences in reproductive energy allocation (15). Further investigation into these factors could provide crucial insights into the mechanisms underlying the timing, duration, and depth of torpor bouts as well as the overall energy balance during hibernation.

Nutrients in foods consumed during the pre-hibernation and hibernation periods could affect hibernation in many hibernators (16, 17). Dietary composition, particularly the balance of polyunsaturated fatty acids (PUFAs), has been proposed to influence the occurrence of torpor and energy expenditure during hibernation (18–20). Early studies have demonstrated that ground squirrels fed diets rich in linoleic acid exhibited longer and deeper torpor bouts (18, 21). Differences in dietary PUFA content could affect both the timing of hibernation onset and depth of torpor across various species (14, 22). For instance, in alpine marmots (*Marmota marmota*), dietary PUFA supplementation affects torpor patterns, with higher levels of n-6 PUFAs associated with longer torpor bouts and lower minimum body temperature (19). In eastern chipmunks (*Tamias striatus*), individuals that exhibited longer torpor bouts possessed higher proportions of n-6 PUFAs in their white adipose tissues (20). The n-6 to n-3 PUFAs ratio may also affect hibernation pattern. In yellow-bellied marmots (*Marmota flaviventris*), a higher n-6: n-3 PUFA ratio in adipose tissue is associated with earlier hibernation onset and longer hibernation duration (22, 23). Garden dormice (*Eliomys quercinus*) fed a diet rich in n-3 PUFAs showed delayed onset of hibernation (24). These lines of evidence suggest that the balance of PUFAs may modulate hibernation timing and patterns across hibernating species. However, changing the dietary n-6: n-3 PUFA ratio did not affect hibernation patterns in arctic ground squirrels (*Urocitellus parryii*) (16). Therefore, the effects of dietary PUFAs on hibernation patterns remain unclear. Cellular PUFA composition may affect membrane fluidity, which is crucial for maintaining cellular function at low temperatures. Additionally, PUFAs may influence the activity of enzymes involved in lipid metabolism, which is critical during hibernation when animals rely primarily on fat stores for energy (25).

In addition, lipophilic antioxidants, such as vitamin E, are crucial for preventing oxidation of PUFAs and may also have an impact on hibernation (26). We previously found that the hepatic amount of α-tocopherol, a natural vitamin E isoform obtained from diets and stored in the liver and other organs via specific transport and binding proteins, affects the resistance to cold-induced cell death of hepatocytes in Syrian hamsters (27, 28). In particular, primary hepatocytes obtained from hamsters fed with the STC diet containing low α-tocopherol amount lost the resistance to cold-induced cell death, whereas those from hamsters fed with the STD diet containing high α-tocopherol amount. This finding using *in vitro* culture system suggests that vitamin E may play a crucial role in protecting tissues from oxidative damage during the extreme temperature fluctuations experienced during hibernation. The potential interaction between PUFAs and lipophilic antioxidants, such as vitamin E, during hibernation is particularly interesting. Although PUFAs are essential for maintaining membrane fluidity at low temperatures, they are highly susceptible to oxidative damage. Vitamin E may help protect these crucial fatty acids from peroxidation during the oxidative stress associated with arousal from torpor (29). However, it remains unclear how distinct diets containing different amounts of vitamin E and PUFAs affect hibernation patterns *in vivo*. The complex interplay between these dietary components and their effects on various aspects of hibernation physiology, including torpor bout duration, minimum body temperature, arousal frequency, and overall hibernation duration, warrant further investigation. Such research could provide valuable insights into the nutritional requirements for successful hibernation, and the mechanisms by which diet influences this remarkable physiological state.

Likewise, there remains room for further investigation of the sex differences in hibernation patterns. Sex differences in hibernation duration have been documented in many species, with females typically hibernating longer than males in arctic ground squirrels (30), alpine marmots (31), Columbian ground squirrels (*Urocitellus columbianus*) (32), golden-mantled ground squirrels (*Callospermophilus lateralis*) (33), and edible dormice (*Glis glis*) (34). For example, female arctic ground squirrels typically emerge from hibernation 10 to 14 days later than do males (30). Interestingly, hamsters appeared to show more complex patterns of sex differences during hibernation. European hamsters (*Cricetus cricetus*) show a reversed pattern in which adult females hibernate for shorter periods than males (17), whereas in Turkish hamsters (*Mesocricetus brandti*), females hibernate longer than males (35). These data suggest that the regulation of sex-specific hibernation patterns in hamsters might be more complicated and less well understood than in other hibernators. However, the extent and mechanisms of these sex-based differences remain poorly understood across hibernating species.

In Syrian hamsters (*Mesocricetus auratus*), there is limited information regarding the sex differences in hibernation patterns. One recent study reported no significant sex differences in hibernation duration, with high mortality rates observed in both sexes, thus complicating the interpretation of results (36). The Syrian hamster is a hibernator that stores and eats food in its burrow during hibernation. This is in striking contrast to fat-storing hibernators, such as ground squirrels, which do not eat food and utilize body fat to survive during the hibernation period (11). As such, hibernation of Syrian hamsters is thought to be more susceptible to dietary influences during hibernation period than that of fat-storing species, as demonstrated in European hamsters (*Cricetus cricetus*) (37). Furthermore, Syrian hamsters are capable of entering torpor after several months of acclimation to environmental cues such as a short photoperiod and cold, unlike obligate or strongly seasonal hibernators that hibernate seasonally under the regulation of intrinsic circannual rhythm (4, 8, 38). These characteristics make Syrian hamsters an ideal model for investigating the potential effects of nutrition on hibernation physiology.

Recent increases in the capacity of data loggers have made it possible to acquire body temperature data for more than a year. This technological advancement has opened new avenues for long-term studies of hibernation patterns in free-living animals (39–41). Previous studies have analyzed long-term Tb datasets using custom-made in-house programs or scripts (24, 40, 42–44). However, lack of standardized, open-source tools has hampered the analysis and comparison of detailed hibernation patterns, including the quantification of each episode of torpor and arousal, the rate of Tb changes, and other important parameters across different studies (8).

In this study, we developed an open-source program, TOHMIN, to facilitate the analysis of long-term Tb data obtained by data loggers (iButton). As a proof-of-concept demonstrating the utility of our analytical tool for detailed hibernation pattern analysis, we investigated whether diet and sex influenced each event of the hibernation cycle in Syrian hamsters.

## 2. Material and methods

### 2.1. Animals, housing, diets, and hibernation induction

Male and female Syrian hamsters were purchased from SLC Inc. to compare the effects of diet and sex on hibernation patterns, male and female Syrian hamsters were purchased from a breeding company (SLC *Inc.*, Japan) at 4 weeks of age, raised in our laboratory under summer-like conditions (L:D cycle = 14:10 h, ambient temperature of 22-25 °C), and subsequently reared under winter-like conditions (L:D cycle = 8:16, ambient temperature of 5 °C) to hibernate as described previously (44). Under winter-like conditions, the animals were housed individually in polypropylene cages to eliminate social interactions. They were provided with ad libitum access to water and food, either an MR standard (STD) diet or an MR stock (STC) diet (Nihon Nosan, Japan). The STD diet contains approximately 5-fold more Vitamin E than the STC diet (27).

### 2.2. Tb recording

Tb recordings with rubber-coated Tb data loggers (iButton, #DS1925L-F5, Maxim Integrated, USA) were surgically implanted into the abdominal cavity of hamsters at 8-10 weeks of age, under anesthesia with 3-4% isoflurane, as previously described (44). All iButtons were programmed to record Tb every 10 min.

### 2.3. Program for analyzing hibernation patterns

The program used to analyze hibernation patterns in this study was developed using Python. It is freely available in the GitHub repository [https://github.com/HIBlab-ILTS/TOHMIN]. Details regarding the program design, functionality, and usage are found in the Results section and in the README file of the GitHub repository.

### 2.4. Statistical analysis

Statistical analyses were performed using Python and R software. The sample sizes used for hibernation period analysis in this study were N = 7 for STD-fed male hamsters, N = 5 for STC-fed male hamsters, and N=5 for female hamsters. Statistical comparisons between the two groups (STD vs. STC in males, and male vs. female) were performed using either Welch’s t-test or the Mann-Whitney U test. The Mann-Whitney U test was used to compare the distributions of PA and DT durations between groups, while Welch’s t-test was applied for other comparisons between two independent groups.

## 3. Results

### 3.1. Design concept and development of the “TOHMIN” program

The “TOHMIN” (named after pronunciation of the Japanese word for hibernation “toumin”) program is a tool for facilitating the analysis of long-term Tb data obtained by data loggers (iButton) from small mammalian hibernators that undergo torpor-arousal cycles during hibernation. This program extracted measurable values for events related to hibernation (Figure 1). The threshold values used to extract the measurable values are presented in the text. For detailed information regarding the threshold values used in the analysis, please refer to our GitHub repository containing the program documentation and source code.

**Figure 1.**
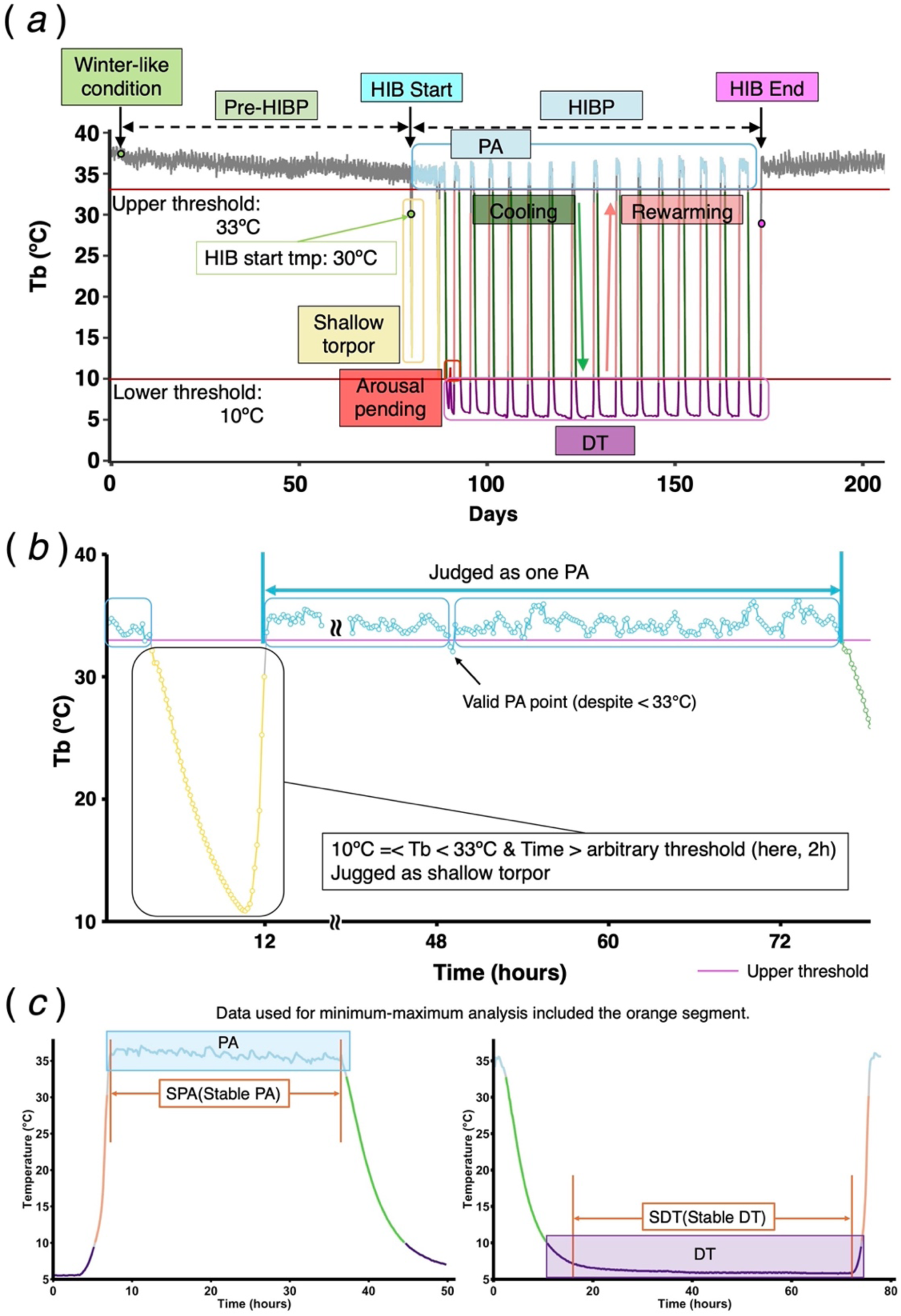
Hibernation pattern analysis and key definitions used in the TOHMIN. (a) Typical changes in body temperature (Tb) of a Syrian hamster exposed to constant winter-like conditions in a laboratory. Key terms are defined as follows: Pre-HIB (pre-hibernation period), HIB Start (hibernation onset), HIB (hibernation period), HIB End (hibernation end date), PA (periodic arousal), and DT (deep torpor). (b) Schematic representation of the program’s strategy to avoid brief drops in Tb during PA-bout, dividing the PA into multiple distinct PAs, an auxiliary threshold based on the duration of a drop was implemented. Double-wave lines indicate the omission of the time-series data in the plot. (c) Illustration of method used to identify stable periodic arousal (SPA) and stable deep torpor (SDT) periods. For the SPA, the starting point was defined as the first plateau reached after rewarming. For SDT, an opposite approach was used to identify the stable period during cooling.

**Hibernation start (HIB start)**: The first time point from which body temperature continues to drop below 30°C for ≥ 1 h (Figure 1*a*).

**Periodic arousal (PA)**: The period during which the body temperature was 33°C or higher after HIB started. The end of each PA episode was defined as the timepoint at which the Tb continued to drop below 33°C for a user-defined duration, which was set to 120 min in this study. This adjustable time criterion was introduced to avoid judging a slight drop in Tb as entering shallow torpor, thereby allowing the two originally divided PA episodes to be regarded as a single episode (Figure 1*b*). A duration of 120 min was chosen based on our preliminary analysis of the temperature patterns, but users can modify this parameter according to their specific research needs and the characteristics of their study species.

**Deep torpor (DT)**: The period during which Tb was below 10°C.

**Shallow torpor (ST)**: The period during which Tb falls below 33°C but does not fall below 10°C.

**Cooling**: The period during which Tb declined from 33°C to 10°C.

**Rewarming**: The period during which Tb increased from 10°C to 33°C. If Tb exceeded 10°C but did not increase to 33°C, it was defined as **Arousal pending**.

**Pre-hibernation period (Pre-HIBP)**: The duration from the day of transition to the cold and short photoperiodic room (the start of the winter-like condition) to the start of HIB.

**Hibernation period (HIBP)**: The duration from HIB start to the end of either spontaneous hibernation end or forced stop of hibernation caused by spontaneous death or experimental death. In this program, the spontaneous end of HIBP was defined as the timepoint after which the Tb did not fall below 33°C for more than 1 h over 2 weeks.

**Death:** If Tb remained below 10°C over 1 week, it was determined as an **Unexplained death** of the animal because, in our experience, Syrian hamsters cannot endure such a long DT state. The point of unexplained death was judged as the date and time when Tb reached a minimum during the period required to determine the death. The program includes a feature to discriminate animal sacrifice for experimental purposes from unexplained death as follows.

**The end of the experiment**, when an animal was sacrificed for experimental purposes, the iButton was removed during dissection, resulting in final temperature readings matching room temperature (20-25°C). The program interprets this as animal sacrifice and not as an unexplained death during hibernation. Alternatively, the end of data can be set manually for the end of the data.

During the initial analysis based on the threshold values for PA and DT, we observed that some PA episodes were divided into two or more segments owing to a transient decrease in Tb below the upper threshold, which impedes accurate counts of PA occurrences. To avoid counting the transient Tb decrease as PA, we implemented a modification to identify stable temperature periods within PA and DT, designated as **Stable PA (SPA)** and **Stable DT (SDT)**, respectively (Figure 1*b, c*). For PA, the SPA period was determined by identifying the plateau phase after the rewarming. The start point of the SPA was defined as the first point at which the body temperature reached a plateau after rewarming towards the PA. A plateau was identified when two consecutive data points showed no increase in temperature (i.e., when the next value was less than or equal to the current value). The endpoint of the SPA was set at the last stable temperature before it began to consistently drop below 33°C. Similarly, for the DT, the SDT period was identified using the opposite approach. Three key temperature metrics were calculated for the SPA and SDT periods as follows: 1. Maximum temperature (max): highest temperature recorded during the stable period. 2. Mean temperature (mean): Average temperature throughout the stable period. 3. Minimum temperature (min): lowest temperature observed during the stable period. This refined approach allows for a more precise analysis of body temperature characteristics during the most stable phases of PA and DT, thereby providing deeper insight into temperature regulation and stability during these critical periods.

The default threshold values were optimized based on our dataset. However, users could adjust these values to optimal values to perform more detailed analyses. This program enables the plotting of time-series data and selection of data for analysis. An advanced feature allows users to selectively exclude periods that contain noise or data-logger errors. The program also facilitates visualization by plotting the analyzed data. The results are displayed for each event in a time series diagram, enabling efficient identification of patterns and anomalies. The following sections present specific examples of the analytical capabilities of the program in various scenarios.

### 3.2. CASE1: Assessment of dietary effects on hibernation patterns

To assess the effects of diet on hibernation, we applied TOHMIN to Tb datasets obtained from hamsters fed two distinct diets: STD or STC diets. Pre-HIBP tended to be longer in STC-fed hamsters than in STD-fed hamsters (p = 0.0576), suggesting that STC-fed hamsters had delayed hibernation onset compared to STD-fed hamsters under winter-like conditions (Table 1).

**Table 1.**
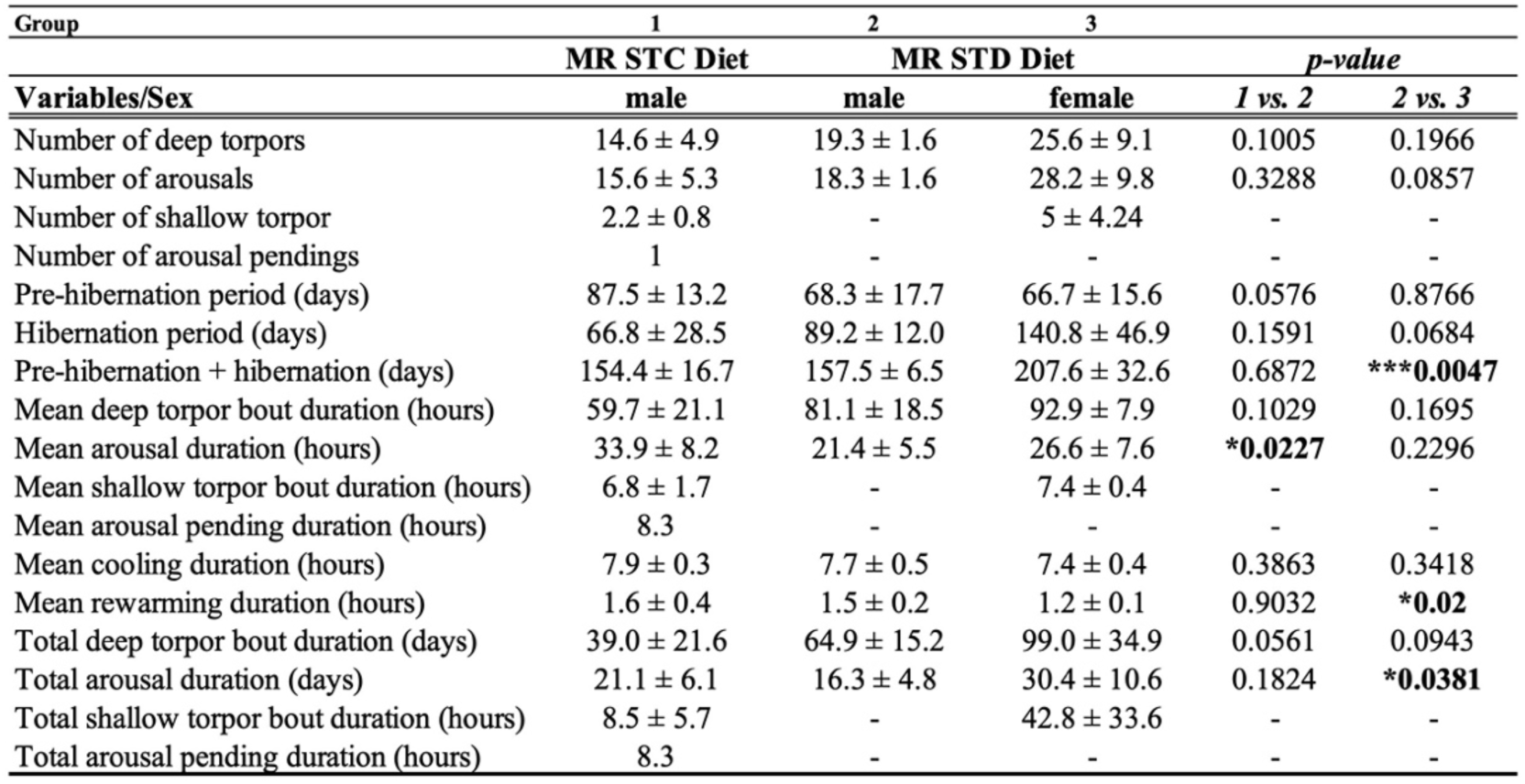
Variability of hibernation patterns in hamsters fed STD or STC diets and male and female. Asterisks indicated statistical differences by t-test. Values indicate mean ± SE.

| Group | 1 | 2 | 3 |  |  |
| --- | --- | --- | --- | --- | --- |
|  | MR STC Diet | MR STD Diet |  | *p-value* |  |
| Variables/Sex | male | male | female | 1 vs. 2 | 2 vs. 3 |
| Number of deep torpor | 14.6 ± 4.9 | 19.3 ± 1.6 | 25.6 ± 9.1 | 0.1005 | 0.1966 |
| Number of arousals | 15.6 ± 5.3 | 18.3 ± 1.6 | 28.2 ± 9.8 | 0.3288 | 0.0857 |
| Number of shallow torpor | 2.2 ± 0.8 | - | 5 ± 4.24 | - | - |
| Number of arousal pendings | 1 | - | - | - | - |
| Pre-hibernation period (days) | 87.5 ± 13.2 | 68.3 ± 17.7 | 66.7 ± 15.6 | 0.0576 | 0.8766 |
| Hibernation period (days) | 66.8 ± 28.5 | 89.2 ± 12.0 | 140.8 ± 46.9 | 0.1591 | 0.0684 |
| Pre-hibernation + hibernation (days) | 154.4 ± 16.7 | 157.5 ± 6.5 | 207.6 ± 32.6 | 0.6872 | *** **0.0047** |
| Mean deep torpor bout duration (hours) | 59.7 ± 21.1 | 81.1 ± 18.5 | 92.9 ± 7.9 | 0.1029 | 0.1695 |
| Mean arousal duration (hours) | 33.9 ± 8.2 | 21.4 ± 5.5 | 26.6 ± 7.6 | **\*0.0227** | 0.2296 |
| Mean shallow torpor bout duration (hours) | 6.8 ± 1.7 | - | 7.4 ± 0.4 | - | - |
| Mean arousal pending duration (hours) | 8.3 | - | - | - | - |
| Mean cooling duration (hours) | 7.9 ± 0.3 | 7.7 ± 0.5 | 7.4 ± 0.4 | 0.3863 | 0.3418 |
| Mean rewarming duration (hours) | 1.6 ± 0.4 | 1.5 ± 0.2 | 1.2 ± 0.1 | 0.9032 | **\*0.02** |
| Total deep torpor bout duration (days) | 39.0 ± 21.6 | 64.9 ± 15.2 | 99.0 ± 34.9 | 0.0561 | 0.0943 |
| Total arousal duration (days) | 21.1 ± 6.1 | 16.3 ± 4.8 | 30.4 ± 10.6 | 0.1824 | **\*0.0381** |
| Total shallow torpor bout duration (hours) | 8.5 ± 5.7 | - | 42.8 ± 33.6 | - | - |
| Total arousal pending duration (hours) | 8.3 | - | - | - | - |

We then evaluated the effects of the different diets on hibernation patterns. Shallow torpor and arousal pending events were detected only in STC hamsters. The mean arousal duration in the STC-fed hamsters was significantly longer than that in the STD-fed hamsters. The cumulative probability distribution of PA durations showed significant differences between diet groups (Figure 2*a*, p = 0.0130), with STC-fed hamsters showing a longer PA duration (median: 21.2 hours) than STD-fed hamsters (median: 17.8 hours). On the other hand, whereas total arousal duration episodes did not differ between the two groups of animals fed different diets, total deep torpor duration tended to be shorter in STC-fed hamsters than in STD-fed hamsters (p = 0.0561). This difference in DT patterns was further supported by the cumulative probability distribution analysis, which showed significant differences between diet groups (Figure 2*c*, p = 0.0005), with STC-fed hamsters having shorter DT duration (median: 70.8 h) than STD-fed hamsters (median: 85.3 hours). No significant differences were found in the mean, maximum, and minimum body temperatures during DT and PA among the different diet groups (Table 2).

**Figure 2.**
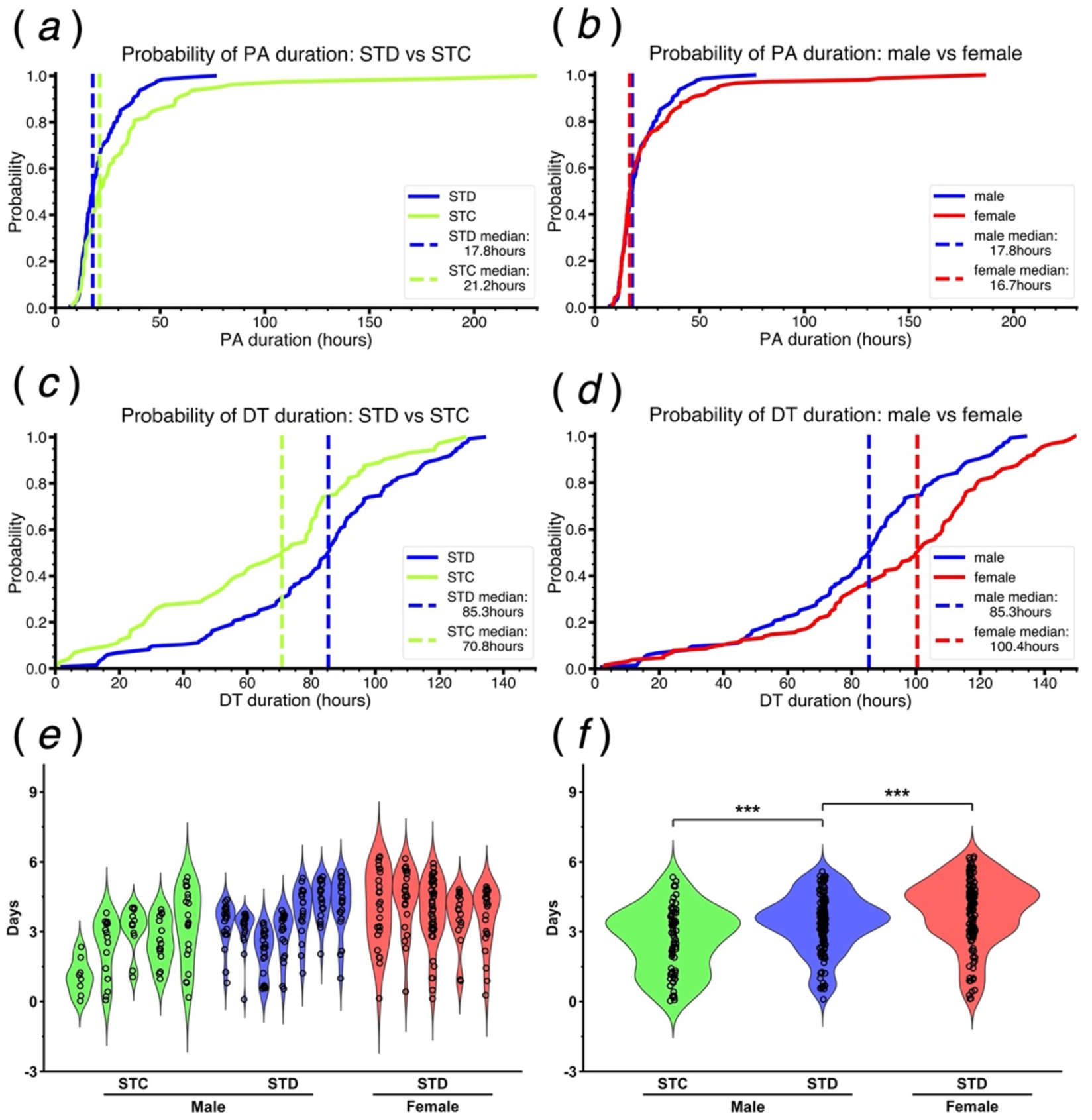
Cumulative probability distribution and violin plots of DT and PA duration in Syrian hamsters. (*a*-*d*) Cumulative probability distributions of PA (*a*, *b*) and DT (*c*, *d*) durations. Comparisons are shown between diet groups (STD: blue lines vs STC: green lines; *a*, *c*) and sexes (males: blue lines vs females: red lines; *b*, *d*). Vertical dashed lines indicate median durations. For PA, medians were STD: 17.8h vs STC: 21.2h (*a*; p = 0.0130) and males: 17.8h vs females: 18.7h (*b*; p = 0.9981). For DT, medians were STD: 85.3h vs STC: 70.8h (*c*; p = 0.0005) and males: 85.3h vs females: 100.4h (*d*; p = 0.0007). Sample sizes were n = 7 for STD males, n = 5 for STC males, and n = 5 for females. All statistical comparisons were performed using Mann-Whitney U tests. (*e*) Each violin plot represents distribution of duration of DT bout per animal, with colours indicating their respective groups (green: STC-fed male, blue: STD-fed male, red: STD-fed female). Black dots represent duration of individual DT bouts. (*f*) Comparison of DT duration among STC-fed male (green), STD-fed male (blue), and STD-fed female (red) groups. The width of each violin represents the probability density of DT durations within group, and black dots indicate duration of DT bouts. Statistical comparisons between groups were performed using Mann-Whitney U tests (***p < 0.001).

**Table 2.** The maximum, minimum, and mean values of body temperature during DT and PA of hamsters. Asterisks indicated statistical differences by t-test. Values indicate mean ± SE.

| Group |  | 1 | 2 | 3 |  |  |
| --- | --- | --- | --- | --- | --- | --- |
|  |  | MR STC Diet | MR STD Diet |  | *p-value* | *p-value* |
| Event | Variables | male | male | female | *1 vs. 2* | *2 vs. 3* |
| DT | mean | 6.0±0.6 | 5.6±0.3 | 5.6±0.2 | 0.2413 | 0.8171 |
|  | min | 5.7±0.6 | 5.4±0.3 | 5.4±0.2 | 0.2550 | 0.9459 |
|  | max | 6.6±0.5 | 6.4±0.3 | 6.5±0.1 | 0.3193 | 0.4104 |
| PA | mean | 35.4±0.2 | 35.4±0.5 | 36.0±0.3 | 0.9373 | **\*0.0364** |
|  | min | 34.0±0.2 | 34.1±0.7 | 34.8±0.3 | 0.6650 | **\*0.0485** |
|  | max | 36.6±0.3 | 36.6±0.4 | 37.0±0.3 | 0.8991 | **\*0.0296** |

### 3.3. CASE2: Comparison of hibernation patterns between male and female hamsters

We analyzed Tb datasets from male and female STD-fed hamsters to examine the sex differences in hibernation patterns. This analysis revealed that the duration of pre-HIBP did not differ between sexes. On the other hand, the HIBP tended to be longer in female than in male (p = 0.0684) (Table 1), and the sum of the pre-HIBP and HIBP was significantly longer in females than in males. These results suggest that female hamsters hibernated for longer than males. In addition, females had significantly longer total PA duration, although the cumulative probability distribution of individual PA durations showed no significant difference between sexes (Figure 2*b*, p = 0.5981; male median: 17.8 hours, female median: 18.7 hours). Although the total and mean DT durations in females did not differ significantly from those of males (Table 2), the cumulative probability distribution of DT durations significantly differed between sexes (Figure 2*d*), suggesting that female hamsters exhibited longer DT durations than males. The median DT duration for females was 100.4 hours, while for males it was 85.33 hours (p = 0.0007). The violin plots further illustrate this sex difference, showing greater variability in DT duration among females than males, with females generally maintaining longer torpor bouts (Figure 2 *e, f*). No significant differences were observed between females and males in the average duration of the cooling, rewarming, DT, and PA phases (Table 2). However, during the PA phase, female hamsters exhibited significantly higher maximum, minimum, and average body temperatures than male hamsters (Table 2). In contrast, during the DT phase, no significant differences were found in the maximum, minimum, or average body temperatures between the sexes (Table 2).

### 3.4. Pre-hibernation period negatively correlates with hibernation duration

Throughout the analysis of Cases 1 and 2, we found a significant negative correlation between the durations of pre-HIBP and HIBP, irrespective of diet and sex (Figure 3). This overall trend suggests that hamsters that took longer to enter hibernation after being transferred to the winter-like condition exhibited shorter HIBP. Consistently, while pre-HIBP tended to be longer in male STC-fed hamsters than in male STD-fed hamsters, the sum of pre-HIBP and HIBP did not differ between the two diet groups of male hamsters (Table 1).

**Figure 3.**
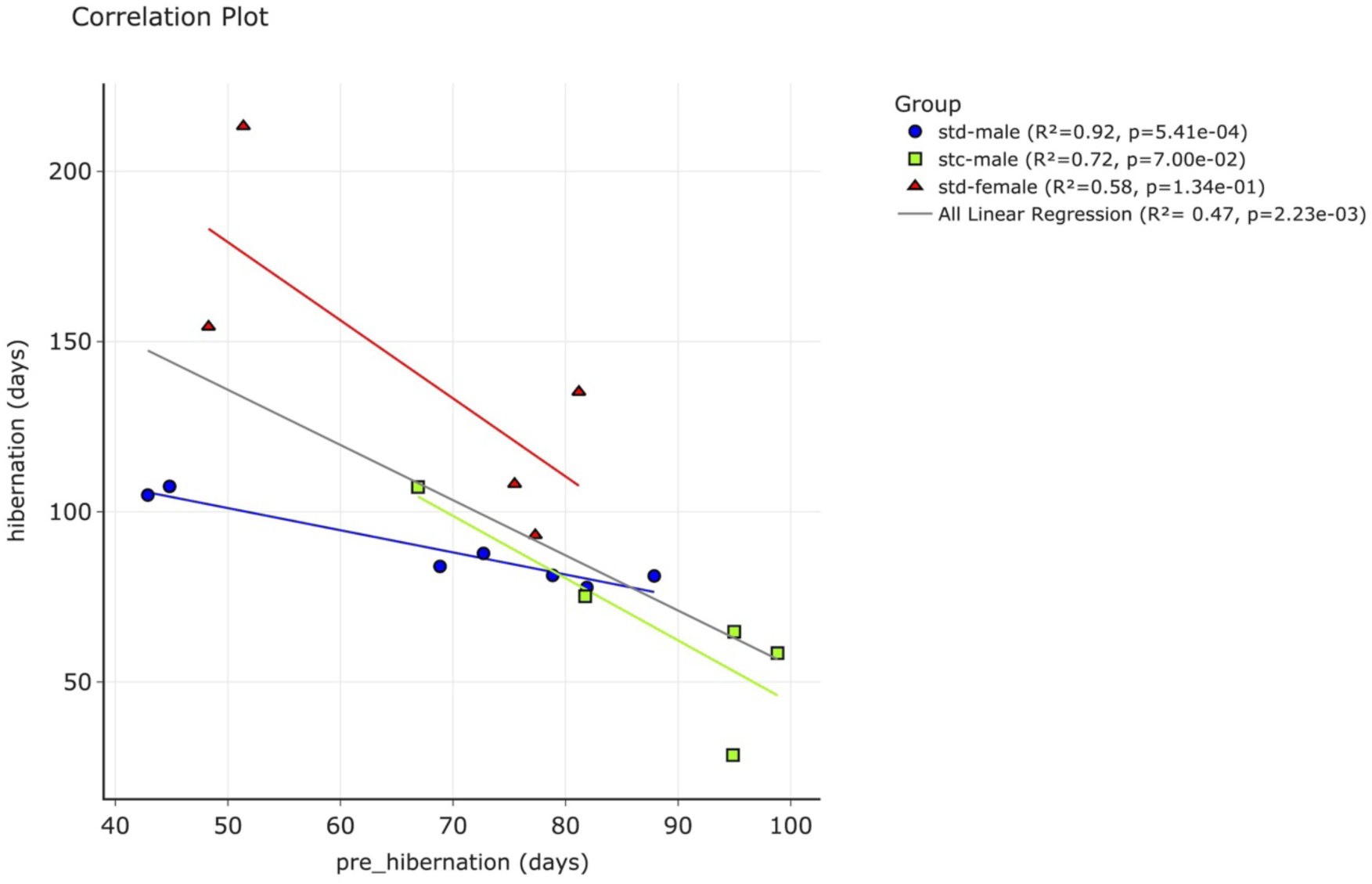
Relationship between hibernation and pre-hibernation periods for all animals. The x-axis represents the pre-hibernation period (days) and the y-axis represents the hibernation period (days). Different colours and symbols represent distinct groups: std-male (blue circles, MRSTD male), stc-male (green squares, MRSTC male), and std-female (red triangles, MRSTD female). The grey line represents a linear regression for all animals, and the coloured line represents a linear regression for each group. The R² values indicate the coefficient of determination for each group and the overall regression. P-values were provided to indicate statistical significance.

## 4. Discussion

In this study, we developed TOHMIN, which enables detailed examination of hibernation patterns. This program employs a threshold-based approach to categorize data into various hibernation states including deep torpor (DT), periodic arousal (PA), shallow torpor (ST), and arousal pending. The ability of the program to handle complex hibernation data through visual reconfirmation and parameter reanalysis addresses a key challenge in hibernation research: the accurate identification and analysis of various physiological states during hibernation (8).

Our analysis revealed significant differences in hibernation patterns between hamsters fed different diets (STD and STC), thereby highlighting the potential influence of nutrition on hibernation physiology. Notably, we observed that STC-fed hamsters exhibited a significantly longer mean arousal duration than did STD-fed hamsters (Table 1). This difference in arousal duration suggests that dietary composition may have a substantial impact on energetics and thermoregulatory processes during PA compared to DT. The observed differences in hibernation patterns between STD and STC diets may be attributed to varying levels of key nutrients, particularly vitamin E and polyunsaturated fatty acids (PUFAs). The STD diet contains approximately 5-fold higher levels of α-tocopherol (vitamin E) than the STC diet (27). Vitamin E is a potent lipophilic antioxidant that plays a crucial role in protecting cellular membranes from oxidative damage, which is particularly important during the rapid metabolic changes associated with arousal from torpor (45, 46).

The higher vitamin E content in the STD diet may contribute to more efficient cellular protection during arousal, potentially allowing shorter periods of arousal. This is consistent with our previous finding that higher hepatic α-tocopherol levels confer increased resistance to cold-induced cell death in Syrian hamsters (27). Our results also support the hypothesis that vitamin E plays a crucial role in protecting against oxidative stress during hibernation and arousal (47). The longer arousal periods observed in STC-fed hamsters may reflect a compensatory mechanism to deal with increased oxidative stress owing to lower antioxidant capacity.

While our study did not directly investigate the effects of PUFAs on hibernation, we observed differences in hibernation patterns between hamsters fed STD and STC diets. Notably, these diets have slightly different PUFA compositions (27), which could potentially contribute to the observed variations in the body temperature patterns during hibernation. Previous research has suggested the potential influence of PUFAs on various aspects of hibernation. For instance, high levels of DHA have been reported to delay hibernation onset in ground squirrels (24), whereas n-3 PUFAs can increase heat production capacity during hibernation (16).

The interplay between PUFAs and dietary antioxidants such as vitamin E in shaping hibernation patterns is complex and warrants further investigation. Future studies should aim to quantify tissue-specific changes in both vitamin E and PUFA levels in response to different diets, and correlate these changes with detailed hibernation parameters. Additionally, examining the expression of genes involved in antioxidant defense and lipid metabolism during different phases of the torpor-arousal cycle could provide mechanistic insights into how dietary components influence hibernation physiology. Thus, our observations suggest that dietary composition, particularly the antioxidant content and fatty acid profile, could significantly influence hibernation patterns in Syrian hamsters. The longer arousal duration observed in STC-fed hamsters may reflect adaptation to a diet with lower antioxidant capacity and different PUFA composition. While these findings highlight the complex relationship between dietary components and hibernation physiology, further research is needed to elucidate the specific roles of vitamin E and PUFAs in the context of hibernation patterns.

Our analysis also revealed significant sex differences in hibernation patterns of Syrian hamsters. Female hamsters typically exhibit longer deep torpor (Figure 2), which is consistent with previous reports of sex differences in torpor duration and HIBP across various hibernator species, including ground squirrels (32, 48–53), alpine marmots (31), and edible dormice (34). Such sex differences during the hibernation period likely reflect different energetic strategies between males and females, possibly related to reproductive demands (54). Females may benefit from longer HIBP to conserve energy for subsequent reproductive efforts, whereas males might emerge earlier to establish territories or compete for mates (33, 55).

However, in the case of hamsters, it is not a simple scenario as to sex influences on hibernation patterns. In Turkish hamsters, females hibernate longer than males, whereas in European hamsters, females hibernate shorter than males (17, 35). In Syrian hamsters, few studies on sex differences in hibernation patterns have been conducted, although one study reported no sex differences in torpor duration in this species (36). However, the hibernation induction and survival rates were very low in this study, hampering interpretation of the results. In contrast, our pilot study using TOHMIN to extract quantitative data on hibernation patterns demonstrated that females hibernate longer than males in this species. Furthermore, the observed differences in body temperature regulation between the sexes, particularly the higher maximum body temperatures in females during arousal, suggest sex-specific thermoregulatory strategies. Such sex-specific differences in thermoregulation may be related to hormonal influences on brown adipose tissue function, which is crucial for heat generation during arousal (56). Further investigation of the physiological mechanisms underlying these sex differences in Syrian hamsters could provide valuable insights into the adaptive significance of sex-specific hibernation strategies.

Another notable finding of our pilot study was the negative correlation between pre-HIBP and subsequent HIBP levels. In other words, individuals who started hibernation later tended to have shorter HIBP, particularly males. Interestingly, in male hamsters, the total duration from the onset of winter-like conditions to the end of hibernation showed a remarkably low individual variation, implying the existence of an intrinsic mechanism that measures this entire period. This consistency might be regulated by seasonal changes in physiological parameters such as reproductive organ size and hormone levels. For instance, in Syrian hamsters, testicular regression occurs during short photoperiod exposure and spontaneous recrudescence begins after a fixed interval (57, 58). These endogenous changes in reproductive physiology may serve as an internal timer that influences the duration of the entire winter period, including both pre-HIBP and HIBP.

In conclusion, TOHMIN facilitates the extraction of many measurable values related to hibernation patterns from long time-series Tb datasets, thereby enabling the precise quantification and detailed analysis of intricate hibernation patterns. Such quantified datasets will lay the groundwork for future investigations into the physiological mechanisms underlying hibernation, the ecological significance of hibernation variability, and potential applications in various fields, ranging from conservation biology to biomedical research.

## Data availability

The datasets utilized and analyzed in this study are available in a previous study (44) and from the corresponding author upon reasonable request. Additional data and information related to this study can be provided upon request.

## Code availability

Data and relevant code for this research work are stored in GitHub: https://github.com/HIBlab-ILTS/TOHMIN and have been archived within the Zenodo repository: https://doi.org/10.5281/zenodo.14327458. The repository contains scripts used for data preprocessing, analysis, and visualization, along with detailed instructions in the README.md file to replicate the results presented in this paper.

## Ethical Compliance

All procedures performed in studies involving animal experiments were done in accordance with the ethical committee of the Hokkaido university, Japan.

## Author Contributions

**Reo Otsuka**: Conceptualization, Data curation, Funding acquisition, Investigation, Methodology, Resources, Software, Validation, Visualization, Writing - original draft, and Writing - review & editing

**Yutaro Shimoyama**: Conceptualization, Data curation, Investigation, Methodology, Software, Validation, Visualization, and Writing - review & editing

**Satoshi Nakagawa**: Data curation, Funding acquisition, Validation, and Writing - review & editing

**Yosifumi Yamaguchi**: (Corresponding Author), Conceptualization, Funding acquisition, Project administration, Resources, Supervision, Validation, Visualization, Writing - original draft, and Writing - review & editing

## Acknowledgments

We would like to thank Sachiyo Enju and Kanako Sone for their experimental assistance, Masamitsu Sone for his advice on data interpretation, and all members of the Yamaguchi Laboratory for helpful discussions and comments.

## Abbreviation

DT: Deep torpor
HIBP: Hibernation period
PA: Periodic arousal
Pre-HIBP: Pre-hibernation period
PUFA: Poly unsaturated fatty acid
SDT: Stable deep torpor
SPA: Stable periodic arousal
ST: Shallow torpor
STD: MR Standard diet
STD: MR Stock diet
Tb: Body temperature
TOHMIN: TOols of Hibernation Measurement and INterpretation

